# Collaboration between the Fab and Fc contribute to maximal protection against SARS-CoV-2 in nonhuman primates following NVX-CoV2373 subunit vaccine with Matrix-M™ vaccination

**DOI:** 10.1101/2021.02.05.429759

**Authors:** Matthew J Gorman, Nita Patel, Mimi Guebre-Xabier, Alex Zhu, Caroline Atyeo, Krista M. Pullen, Carolin Loos, Yenny Goez-Gazi, Ricardo Carrion, Jing-Hui Tian, Dansu Yaun, Kathryn Bowman, Bin Zhou, Sonia Maciejewski, Marisa E. McGrath, James Logue, Matthew B. Frieman, David Montefiori, Colin Mann, Sharon Schendel, Fatima Amanat, Florian Krammer, Erica Ollmann Saphire, Douglas Lauffenburger, Ann M. Greene, Alyse D. Portnoff, Michael J. Massare, Larry Ellingsworth, Gregory Glenn, Gale Smith, Galit Alter

**Affiliations:** Ragon Institute of MGH, MIT, and Harvard, Cambridge, MA 02139, USA; Novavax, Inc., 21 Firstfield Road, Gaithersburg, MD 20878, USA; Department of Biological Engineering, Massachusetts Institute of Technology, Cambridge, MA 02139, USA; Texas Biomedical Research Institute. 8715 West Military Drive, San Antonio, TX 78227, USA; University of Maryland, School of Medicine, 685 West Baltimore St, Baltimore, MD 21201, USA; Department of Surgery, Duke University Medical Center, Durham, NC 27710, USA; La Jolla Institute for Immunology, La Jolla, CA 92037, USA; Department of Microbiology, Icahn School of Medicine at Mount Sinai, New York, NY, USA

**Keywords:** NVX-CoV2373 vaccine, Matrix-M™ adjuvant, SARS-CoV-2 spike glycoprotein, non-human primate, COVID-19

## Abstract

Recently approved vaccines have already shown remarkable protection in limiting SARS-CoV-2 associated disease. However, immunologic mechanism(s) of protection, as well as how boosting alters immunity to wildtype and newly emerging strains, remain incompletely understood. Here we deeply profiled the humoral immune response in a cohort of non-human primates immunized with a stable recombinant full-length SARS-CoV-2 spike (S) glycoprotein (NVX-CoV2373) at two dose levels, administered as a single or two-dose regimen with a saponin-based adjuvant Matrix-M™. While antigen dose had some effect on Fc-effector profiles, both antigen dose and boosting significantly altered overall titers, neutralization and Fc-effector profiles, driving unique vaccine-induced antibody fingerprints. Combined differences in antibody effector functions and neutralization were strongly associated with distinct levels of protection in the upper and lower respiratory tract, pointing to the presence of combined, but distinct, compartment-specific neutralization and Fc-mechanisms as key determinants of protective immunity against infection. Moreover, NVX-CoV2373 elicited antibodies functionally target emerging SARS-CoV-2 variants, collectively pointing to the critical collaborative role for Fab and Fc in driving maximal protection against SARS-CoV-2. Collectively, the data presented here suggest that a single dose may prevent disease, but that two doses may be essential to block further transmission of SARS-CoV-2 and emerging variants.

**Highlights:**

- NVX-CoV2373 subunit vaccine elicits receptor blocking, virus neutralizing antibodies, and Fc-effector functional antibodies.
- The vaccine protects against respiratory tract infection and virus shedding in non-human primates (NHPs).
- Both neutralizing and Fc-effector functions contribute to protection, potentially through different mechanisms in the upper and lower respiratory tract.
- Both macaque and human vaccine-induced antibodies exhibit altered Fc-receptor binding to emerging mutants.

## Introduction

SARS-CoV-2 causes a spectrum of respiratory disease from asymptomatic to mild and severe coronavirus disease (COVID-19). Since it crossed into humans, the virus has spread globally with over 90 million confirmed cases and over 2 million deaths^1^. COVID-19 manifests with a range of clinical symptoms from asymptomatic to severe disease, with 50-75% of infected individuals exhibiting asymptomatic infection and only a small proportion (2-5%) developing severe disease, requiring mechanical ventilation^2-4^. The vaccines authorized for emergency use, mRNA-1273 and BNT162b2, have been successful in preventing severe infections and inducing anti-SARS-CoV-2 CD4+ T cell, CD8+ T cell, and potent neutralizing antibody responses^5-7^. However, whether these vaccines confer protection against transmission as well as disease remains unclear.

Emerging Phase 3 data suggest that vaccine-mediated protection emerges as early as 10 days following primary vaccination^8,9^, at a time when neutralizing antibodies are low or undetectable^5-7^. Similarly, emerging correlates of immunity following administration of DNA- and adenoviral-vector SARS-CoV-2 vaccination point to a potential additional role for added antibody effector functions, in collaboration with neutralization, as key correlates of immunity against SARS-CoV-2^10,11^. However, whether these responses evolve following the prime or the boost, provide differential protection across the upper and lower respiratory tract, and provide protection against variants remains unclear.

In this study, we deeply interrogated humoral correlates of protection in a cohort of rhesus macaques immunized with one or two doses 5 or 25 µg of a stabilized recombinant full-length SARS-CoV-2 spike (S) glycoprotein (NVX-CoV2373) with 50 µg Matrix-M adjuvant. Animals immunized with the two-dose regimen, regardless if given the high (25μg) or low (5μg) antigen dose, were protected against upper and lower respiratory infection (URTI and LRTI) and shedding of replicating virus, while a single vaccine injection (regardless of antigen dose) was only partially protective against infection. Distinct combinations of Fc-features and neutralizing antibody responses were associated with protection in the upper and lower respiratory tract, pointing to potential mechanistic differences required to control the virus at these distinct immunological locations. Critically, the NVX-CoV2373 generated binding and functional humoral immune responses to several emerging SARS-CoV-2 variants. These data point to boosting-driven functional maturation of the humoral immune response as a key immune event required to achieve full protection against infection and transmission of SARS-CoV-2 and emerging mutants.

## Results

### Subgenomic virus mRNA in respiratory samples

Emerging Phase 3 data from mRNA vaccine platforms suggest that vaccine-induced protection against disease is observable as early as 10 days following vaccine priming, prior to the presence of robust neutralizing antibody levels^8,9^. However, whether these responses are associated with complete sterilizing immunity remains unclear. To define the specific humoral profiles that track with protective immunity against disease and infection, we profiled the humoral immune response induced by a stabilized, full-length SARS-CoV-2 Spike (S) vaccine (NVX-CoV2373) following a prime-only or prime/boost vaccine regimen administered at 2 different antigen doses (5 and 25μg) with Matrix-M adjuvant (50μg). Groups of rhesus macaques (n=5) were immunized with one vaccine dose (study day 0) or two vaccine doses, spaced 3 weeks apart (study day 0 and 21). Control animals (n=4) received one or two injections of formulation buffer (placebo). Serum was collected prior to immunization (day 0) and 21 and 31/32 days after the first dose (**Fig 1A**).

**Figure 1.**
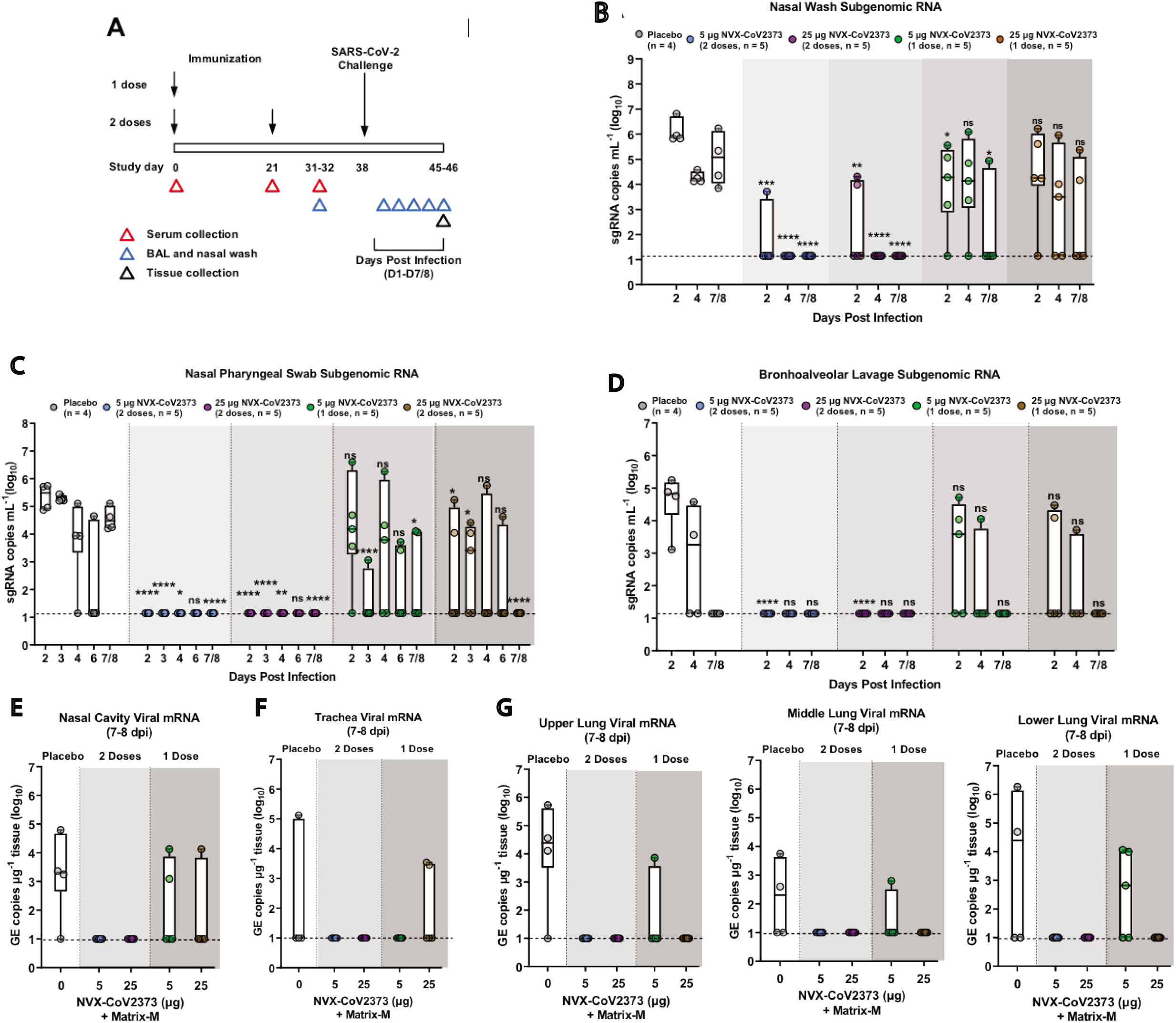
Subgenomic RNA and viral RNA in upper and lower respiratory tract of NVX-CoV2373 immunized rhesus macaques. (**A**) Groups of adult rhesus macaques (n= 4-5/group) were immunized with a single priming dose (study day 0) or a prime/boost regimen (study day 0 and 21) of 5 μg or 25 μg NVX-CoV2373 with 50 μg Matrix-M (0.5mL; IM). A separate group (n=4) received formulation buffer (placebo). Immunized and placebo animals were transferred to an ABSL-3 containment facility (study day 31/32) and acclimated for 7 days prior to challenge with a total of 1.05 × 10^6^ pfu SARS-CoV-2 (USA-WA1/2020 isolate) in 500 μL divided between the intranasal (IN) and intra-tracheal (IT) routes. Animals were monitored daily for up to 7-8 days post infection (1-8 dpi). Serum sample collection days are indicated by the red triangles. Bronchoalveolar lavage (BAL) sample collection days are indicated by the blue triangles. Necropsy and tissue collection is indicated by the black triangle. Quantitative RT-PCR was used to measure the replicating subgenomic (sg) envelope (E) RNA in nasal washes, nasal pharyngeal swabs, and BAL samples collected for up to 7-8 dpi. (**B**) Nasal pharyngeal washes. (**C**) Nasal swabs. (**D**) BAL aspirates. (**E**) SARS-CoV-2 gRNA in the Nasal cavity virus load. (**F**) Trachea virus load. **(G**) Upper, middle and lower lobes of the lungs of immunized and placebo treated animals. In the bar-and-whisker plots, the median is indicated by a horizontal line, the top and bottom of the box indicated the interquartile range, and the whiskers indicate the minimum and maximum values. Individual animal values are indicated by the colored symbols. Dashed horizontal line indicates the limit of detection. Genomic equivalent copies (GE copies mL^-1^). Significant differences between the placebo group and the immunized groups was determined by Student’s t-test (two tailed, unpaired). Not significant (ns), ∗p ≤ 0.05, ∗∗p ≤ 0.01, ∗∗∗p ≤ 0.001, ∗∗∗∗p ≤ 0.0001.

Protection was assessed by analyzing viral loads across the upper (nasal washes and nasal pharyngeal swabs) and lower (bronchoalveolar lavage; BAL) respiratory tract on days 2-8 post-infection (dpi). The highest levels of viral subgenomic RNA (sgRNA) were observed in placebo animals across the upper and lower-respiratory tract samples, with peak viral loads observed 2dpi and persistent sgRNA until day 7/8 (**Fig 1B, C, D**). Animals immunized with a single dose of 5μg or 25μg NVX-CoV2373 had lower levels of replicating virus at day 2 in all tissues compared to placebo, however the 25μg dose was able to clear sgRNA in BAL and nasal pharyngeal swabs at day 7/8, while the 5μg only cleared BAL. The animals that received 5μg or 25μg antigen in a prime/boost regimen had no detectable viral loads in BAL or nasal pharyngeal swabs at any day and all sgRNA was cleared in nasal washes by day 4. In addition, tissue samples were collected from the upper, middle, and lower right lung lobes; trachea; and nasal cavity at the scheduled necropsy (7-8 dpi) and analyzed for viral gRNA. There was no gRNA in nasal cavity, trachea, or lungs of animals immunized with 5μg or 25μg antigen in a prime/boost regimen (**Fig 1E, F, G**). Conversely, nearly all placebo animals exhibited gRNA in each tissue (**Fig 1E, F, G**). Animals immunized with a single vaccine dose were partially protected, with a minority of animals having detectable gRNA. These data suggest that one vaccine dose was able to induce a partially protective immune response, differing by antigen dose level, but two vaccine doses resulted in full protection against infection along the respiratory tract, independent of antigen dose.

### Antibody responses after NVX-CoV2373 immunization

To determine if the humoral immune response could distinguish protected from non-protected animals, we analyzed the IgG titers and neutralizing antibody response across the vaccine groups. Robust anti-S IgG titers were observed across both vaccine groups after a single immunization. Anti-S IgG titers remained stable at 31/32 days after 1 dose, however anti-S IgG titers significantly increased 21-35-fold within 10 days following the booster immunization with 5μg or 25μg of NVX-CoV2373 (**Fig 2A**). Low levels of mucosal anti-S IgG antibodies were detected in the nasal washes and BAL aspirates collected 31/32 days after one immunization, increasing 8-22-fold in nasal washes and BAL aspirates at 10 days following the booster immunization (**Fig 2B, 2C**).

**Figure 2.**
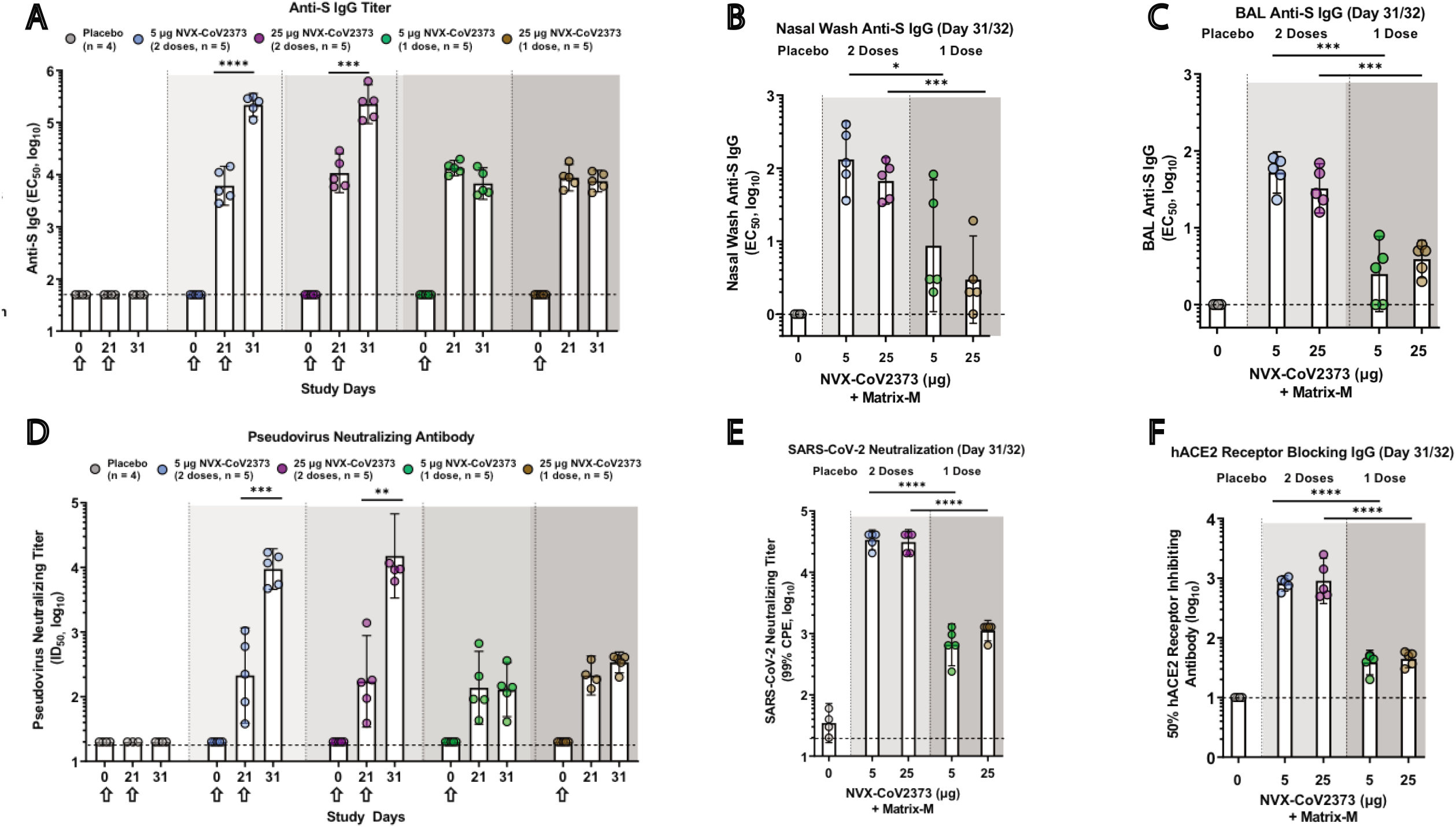
Immunogenicity of NVX-CoV2373 vaccine in rhesus macaques. (**A**) Serum anti-S IgG titer (**B**) Nasal wash and (**C**) bronchoalveolar lavage (BAL) samples were collected 31/32 days after the first immunization and prior to challenge and analyzed for spike (S)-specific mucosal IgG (n= 4-5/group). (**D**) Pseudovirus neutralizing titer (ID_50_). (**E**) SARS-CoV-2 neutralizing antibody titer (99% inhibition of cytopathic effect, 99% CPE) study day 31/32. (**F**) hACE2 receptor blocking antibody titer (study day 31/32). The geometric mean titers (GMT) are indicated by the white bars. Hollow arrows indicate prime/boosting with NVX-CoV2373. The error bars indicate the 95% confidence interval (95% CI). Individual animal values are indicated by colored symbols. A Student’s t-test (unpaired, two tail) was used to compare antibody levels between groups immunized with one and two doses. ∗∗p ≤ 0.001, ∗∗∗p ≤ 0.0001, ∗∗∗∗p ≤ 0.00001. The horizontal dashed line indicates the limit of detection (LOD) for each assay.

To further profile the functional potential of the vaccine induced antibodies, a spike-pseudotype virus neutralization assay was used to assess the neutralizing capacity in serum of immunized animals. Serum from animals immunized with 5μg or 25μg NVX-CoV2373 had similar pseudovirus neutralizing titers (ID_50_) after a single dose. Following the booster immunization, pseudovirus neutralizing titers significantly increased, with no significant differences noted between the antigen doses (**Fig 2D**). In addition, live wild type virus neutralization assays and hACE2 inhibition exhibited similar trends, with detectable neutralization/inhibition at day 21 in all regimens, with a significant increase after the second vaccine dose (**Fig 2E and 2F**). Overall, these results indicate that NVX-CoV2373 administered as a prime/boost regimen elicited high anti-S IgG titers, capable of blocking binding to the hACE2 receptor and neutralizing in vitro infectivity of spike-pseudotyped virus and wild type SARS-CoV-2. All non-human primates (NHPs) treated with one dose had similar neutralization titers, but only some were protected from viral infection, suggesting that neutralization may not be sufficient to fully explain complete protection from infection, particularly following a single vaccine dose.

### System serology profiling

Natural SARS-CoV-2 infection is marked by a rapid rise of multiple antibody isotypes and subclasses, each positioned to recruit a diverse set of antibody effector functions^12,13^. Recent studies have noted a significant correlation between antibody-effector function, rather than neutralization, with natural resolution of infection in humans^14^. Thus, we next examined the evolution of subclass, isotype, Fc-receptor, and Fc-effector function across doses and boosting strategies (**Supplementary Fig 1**).

As expected, based on titers (**Fig 2**), luminex IgG1 levels were robustly induced following a single vaccine dose, indistinguishably across antigen levels, with a 1.5-4-fold increase following a boost (**Fig 3A**). Similarly, IgA were induced robustly to a maximal level after one 25μg dose, but required boosting to reach maximal levels in the 5μg vaccine group (**Fig 3A**). Conversely, a trend towards higher levels of IgM were noted in 5μg vaccine group following a single vaccine dose, that declined with a boost and were largely lost in the 25μg dose group (**Fig 3A**), pointing to enhanced class switching to more mature antibody subclasses with boosting and higher antigen doses. These data point to the first differences across antigen-dosing group, highlighting equivalent IgG and IgA selection across groups, but more aggressive switching of IgM, shifting the polyclonal balance of the vaccine-specific antibody pool towards a more mature Fc-functional profile.

**Figure 3.**
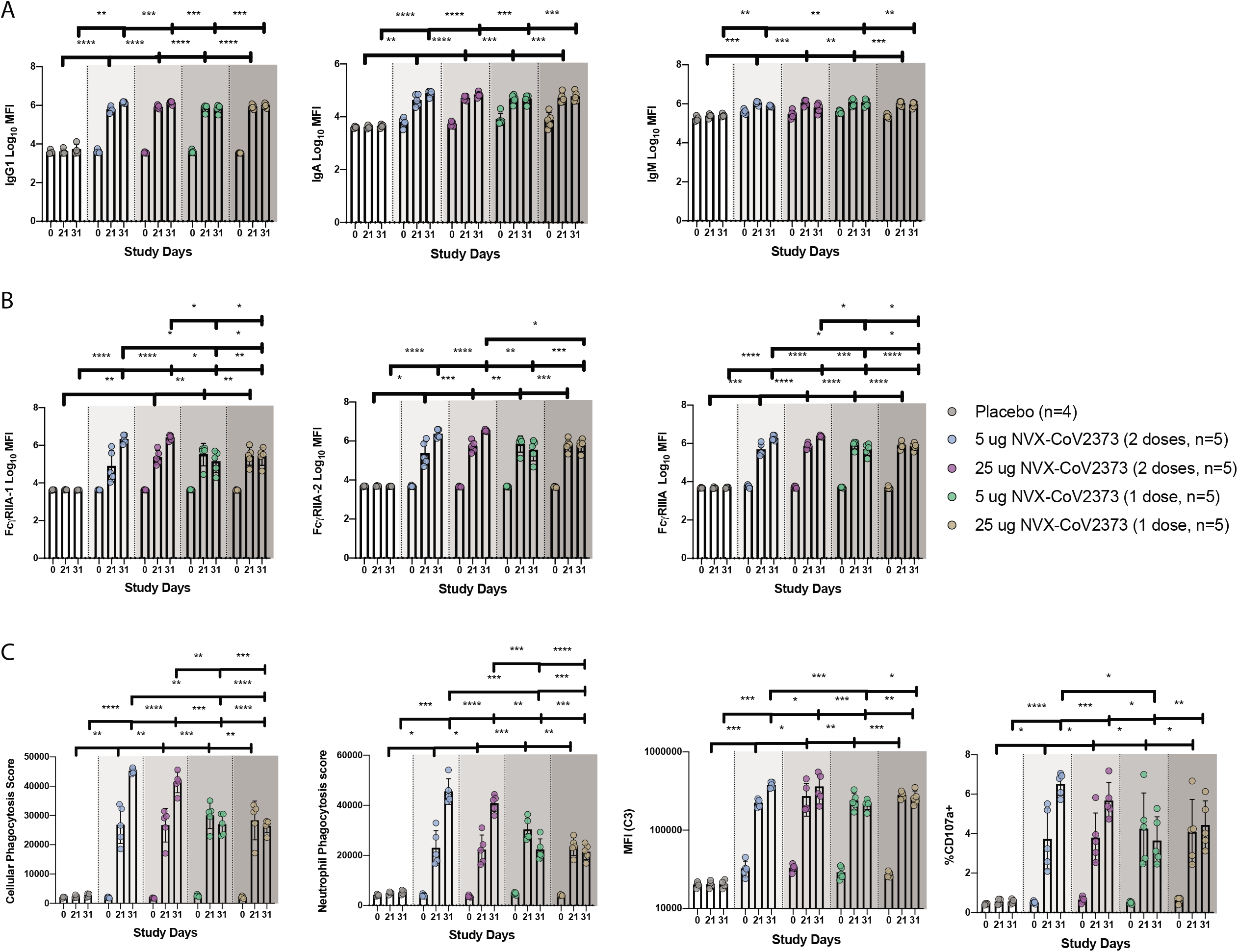
System serology profiling of NVX-CoV2373 immunized rhesus macaques. Serum was collected day 21 and day 31/32 after the first dose of NVX-CoV2373, and profiled for the anti-NVX-CoV2373 antibody response. Luminex was used to quantify the (**A**) antibody isotypes (IgG1, IgA, and IgM) and (**B**) FcR binding (FcγRIIA-1, FcγRIIA-2, FcγRIIIA) for the anti-NVX-CoV2373 antibody response. (**C**) The functional anti-NVX-CoV2373-specific antibody responses for antibody-dependent cellular phagocytosis, antibody-dependent neutrophil phagocytosis, antibody-dependent complement deposition, and antibody-dependent NK degranulation (measured by CD107%). A two-way ANOVA with Tukey correction for multiple comparison was used to compare antibody levels between groups. Not significant (ns), ∗p ≤ 0.05, ∗∗p ≤ 0.01, ∗∗∗p ≤ 0.001, ∗∗∗∗p ≤ 0.0001.

Changes in polyclonal antibody profiles result in the potential formation of distinct swarms of antibodies able to engage with a target pathogen, forming qualitatively distinct immune complexes, that collectively shape the Fc-receptors (FcRs) bound on innate immune cells, thereby driving distinct antibody effector functions^15-18^. Thus, to explore differences in functionality across doses and boosting regimens, we next profiled differences in binding profiles across rhesus Fc-receptors. Equivalent FcγRIIA-1 binding was observed across the 2 antigen doses after the prime, although there was a trend to a loss of binding at day 31/32 in the 5μg dosing group (**Fig 3B**). However, after a boost, FcγRIIA-1 binding antibodies increased by 4-100-fold across the doses, with a trend towards higher binding antibodies in the 25μg dosing group (**Fig 3B**). Nearly identical profiles were observed across the other rhesus FcRs, pointing to a substantial quantitative advantage induced by the boost, that tended to differ across the doses. Finally, to explore the functional impact of these changes in vaccine induced antibody Fc-profiles, we examined the ability of the humoral response to stimulate antibody-dependent functions: cellular monocyte phagocytosis (ADCP), neutrophil phagocytosis (ADNP), complement deposition (ADCD), and NK degranulation (NKdegran). Similar ADCP responses were induced across the antigen doses following a single vaccination (**Fig 3C**). Conversely, robust augmentation of ADCP was observed with a boost (**Fig 3C**), that surprisingly tended to be higher in the 5μg group. An identical profile was observed for NK-cell activating antibodies. Neutrophil phagocytosis was slightly higher in the 5μg group after the prime, and then fully matured across both groups with a boost, remaining slightly elevated in the 5μg group. Conversely, complement activating antibodies were induced equivalently across the antigen-dosing groups following a single dose, and increased with a boost in an antigen dose-independent manner. Thus, while titers and neutralization reached near maximal potential after a single vaccine dose, these data point to a critical role for boosting in driving the full maturation of the Fc-effector potential of the vaccine induced humoral response, that are further subtly tuned by antigen dosing.

### Unique humoral profiles of vaccine regimen

Given the various univariate profile differences noted across the vaccine groups, we next aimed to define whether distinct multivariate profiles were induced across the regimens. Aggregate data clearly highlighted the striking influence of the boost and the more nuanced effects of antigen dose on shaping the polyclonal vaccine response (**Fig 4A**). Antigen-dose effects emerged upon unsupervised analysis using a principal component analysis (PCA), pointing to a tendency towards separation between antigen dose and vaccine-specific antibody profiles in the animals that received a single dose (**Fig 4B**), that was largely lost with the boost (**Fig 4C**). However, integration of the 4 groups clearly demonstrated the dominant influence of the boost in shaping antibody profiles(**Fig 4D**). Specifically, robust separation in antibody profiles across single and double immunized animal vaccine-specific antibody profiles (**Fig 4D**), with a more subtle effect of dose on shaping vaccine-specific antibody profiles, solely observed in the single dose arms. Finally, radar plots of the humoral immune response across vaccine arms demonstrated the clear explosion of humoral immune maturation with the second dose, albeit slight differences in antibody effector functions were noted across the doses. Additionally, more nuanced differences were observed in the single dose arms, with a more balanced functional response observed in the 25μg group compared to the 5μg immunized animals at day 31-32, prior to challenge (**Fig 4E**). These data provide a deep immunologic view of the vaccine-induced polyclonal functional profiles induced following vaccination, and how they are shaped by dose and boosting prior to challenge.

**Figure 4.**
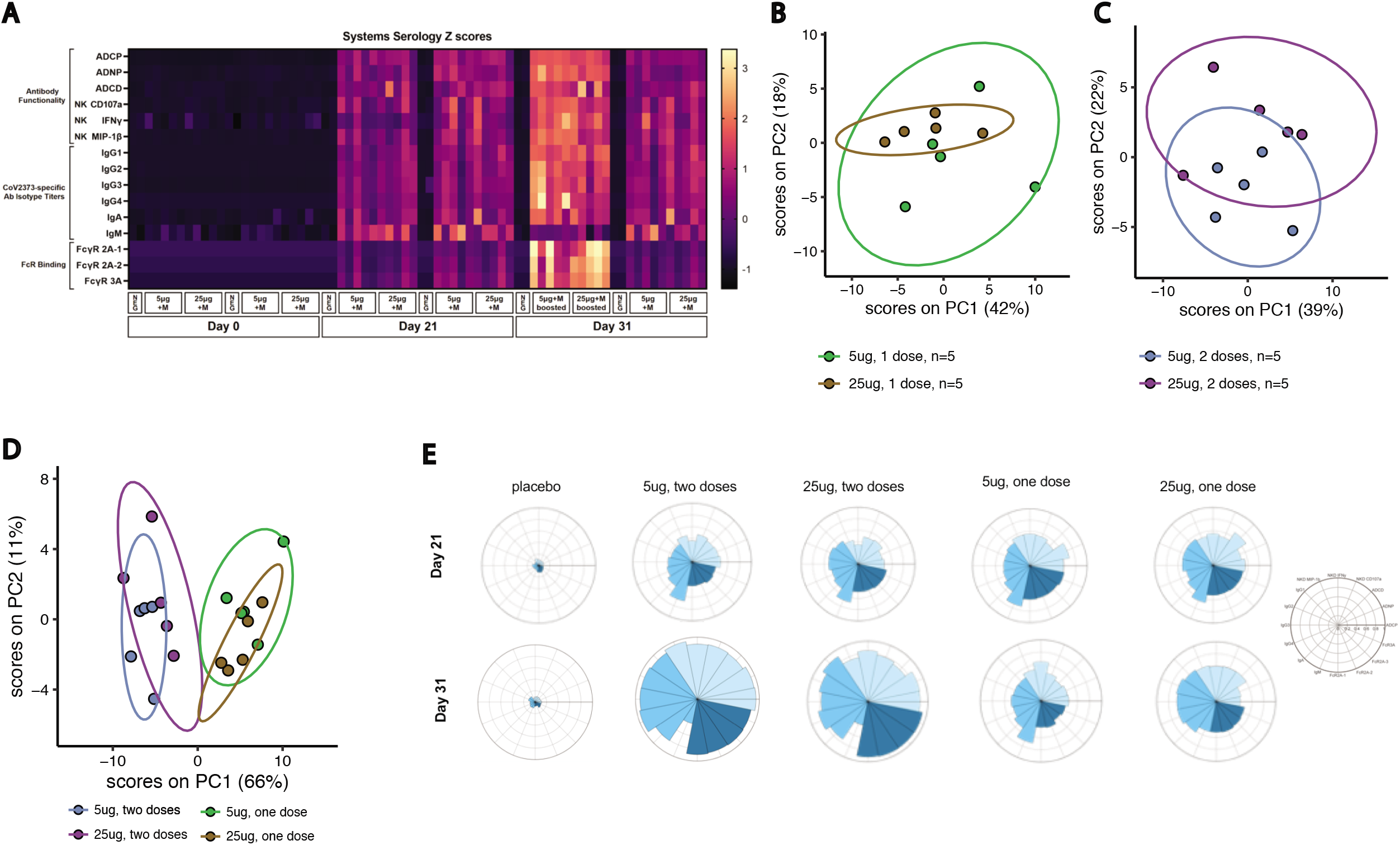
Unique humoral profile of vaccine regimens. Multivariate analysis was performed to distinguish the humoral response between the various vaccine regimens. (**A**) Heatmap of the humoral response to SARS-CoV-2 spike. Each Column is one NHP and one time point. Each row was Z-scored across itself for the whole cohort. (**B**) Principal component analysis (PCA) of antibody features at day 31/32 showing NHPs that received one 5μg dose (light blue) or one 25μg dose (dark blue). Ellipses indicate 90% confidence regions assuming a multivariate t distribution. (**C**) PCA of antibody features at day 31/32 showing NHPs that received two 5μg doses (light pink) or two 25μg doses (dark pink). (**D**) PCA of antibody features at day 31/32 showing NHPs that received one 5μg dose (light pink), one 25μg dose (dark pink), two 5μg doses (light blue), or two 25μg doses (dark blue). (**E**) The radar plots show the median percentile for antibody titer, FcR binding, and antibody function (legend on right) for NHPs treated with placebo, two 5μg doses, two 25μg doses, one 5μg dose, or one 25μg dose in serum collected on day 21 (top row) and day 31/32 (bottom row).

### Immune correlates of protection from viral infection

While neutralizing antibodies have been clearly linked to vaccine-mediated protection following DNA^11^, AD26 ^10^, protein^19^, and mRNA based vaccination ^5-7^, protection has been noted in humans prior to the evolution of neutralizing antibodies^8,9^. Similarly, despite robust induction of neutralizing antibodies given one or two doses of NVX-CoV2373, variable levels of protection were observed against upper and lower respiratory viral loads across the groups (**Fig 1B,C,D,E,F,G**). To define the humoral correlates of immunity of viral control across the respiratory tract, all antibody metrics were integrated, and an unsupervised multivariate analysis was performed to objectively define antibody correlates of immunity. Clear separation was noted in vaccine-induced antibody profiles across NHPs exhibiting complete protection against SARS-CoV-2 compared to animals that exhibited viral loads in one or several compartments (**Fig 5A**). Specifically, the PCA illustrated a substantial split in antibody profiles in animals that exhibited no protection/protection in the lower respiratory tract (BAL) from animals that exhibited more complete protection across the upper and lower-respiratory tract (nasal washes, nasal swabs, and BAL). Thus, unsupervised analysis suggested the presence of unique humoral immune correlates of immunity in lower and upper respiratory tracts.

**Figure 5.**
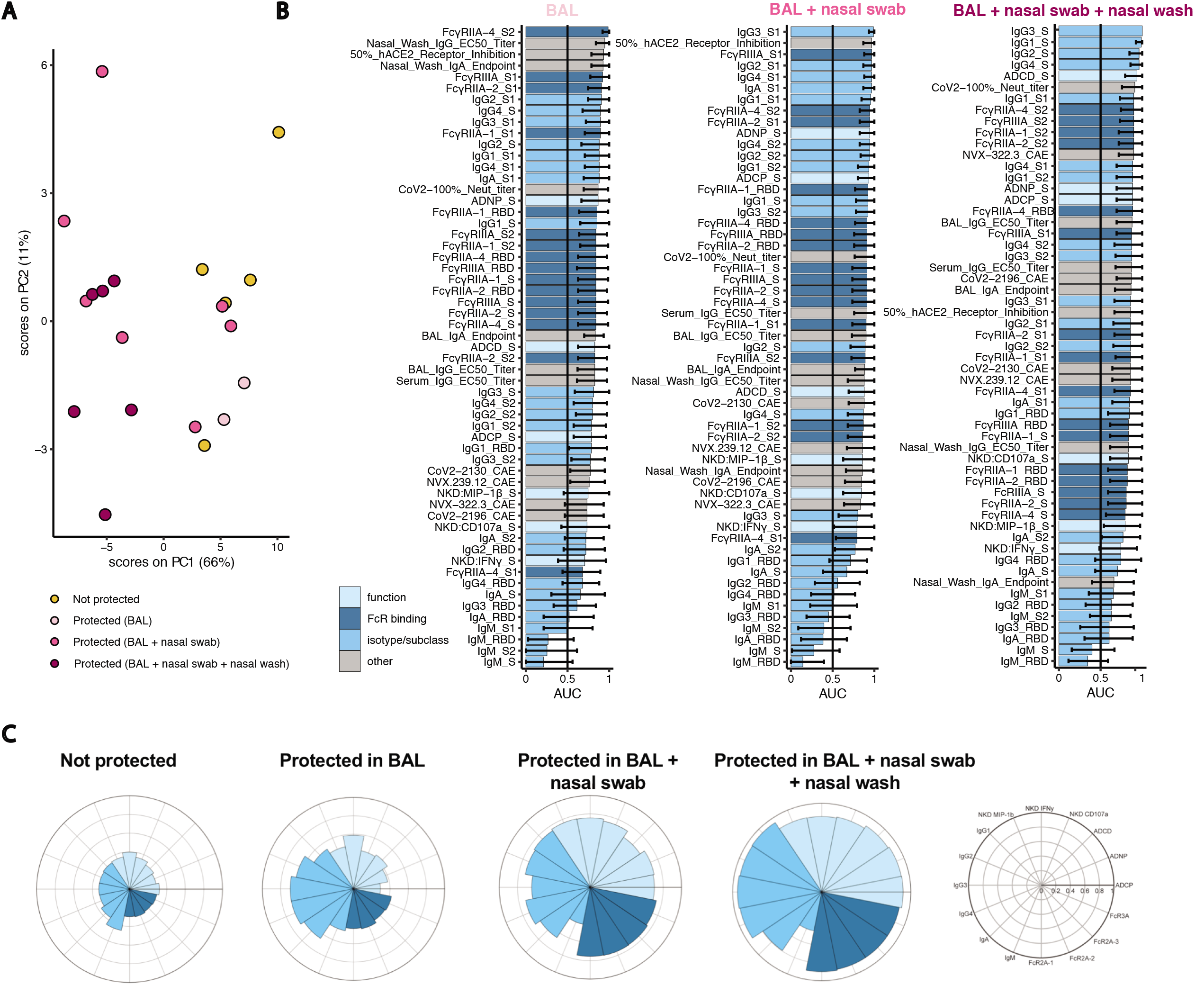
Immune correlates of protection from viral replication. Multivariate analysis was performed to identify the features of a protective humoral response. (**A**) Principal component analysis (PCA) for the immunized NHPs (n=20, no placebos included) indicating protected (blue) NHPs with no detectable virus in BAL, BAL + nasal swab, BAL + nasal swab + nasal wash vs non-protected (yellow) NHPs. Ellipses indicate 90% confidence regions assuming a multivariate t distribution and are shown for protected and non-protected NHPs. (**B**) Correlates of protection for BAL (n=20), nasal swab (n=20), or nasal wash (n=19) at day 31/32. The area-under-the-curve (AUC) for the receiver operator characteristic (ROC) curve is shown with 95% confidence intervals for each antibody feature. (**C**) The radar plots show the median percentile for antibody titer, FcR binding, and antibody function (legend on right) for non-protected, protected in BAL, protected in BAL+nasal swab, or protected in BAL+nasal swab+nasal wash NHPs.

To gain deeper resolution into the specific features of the humoral immune response that may lead to these distinct levels of viral restriction across compartments, the relationship of individual features and protection was assessed by calculating the area-under-the-curve for each receiver operator characteristic (ROC) curve within each compartment (**Fig 5B**). The top features associated with protection in the lower respiratory tract (BAL) included antibody titers, S2- and S1-specific FcR binding, and hACE2 receptor inhibition. Similarly, the top features associated with protection in the BAL and nasal pharyngeal swab included the levels of S1-specific antibody titers of several IgG subclasses and hACE2 inhibition. However, complete protection from viral replication across the upper and lower respiratory tracts was associated with a robust whole S-specific multi-subclass specific response, complement-depositing functions, and neutralizing antibody titers. The radar plots further illustrated the magnitude and multivariate nature of the protective humoral immune response, marked by poor antibody responses in unprotected animals, an expansion of subclasses, but not functions, in animals with solely lower respiratory tract protection (BAL), an expanded functional and FcR-binding antibody profiles in animals with BAL and nasal swab protection. Conversely, the largest, functionally expanded humoral immune response was observed in animals with complete protection across the upper and lower respiratory tract (**Fig 5C**). These data point to an intimate collaboration between the Fc and Fab in driving full viral protection, where neutralization may be key to lower-respiratory protection, but the potential need for additional Fc-effector functions in collaboration with neutralization may be key for full protection across the respiratory tract.

### Antibody response to emerging SARS-CoV-2 mutants

Despite the promising results observed in Phase 3 trials with the mRNA vaccines, significant concern has recently arisen globally, due to the rapid emergence of SARS-CoV-2 variants across the globe^20-23^. Among the variants, the recently identified genetic variant, B.1.1.7 (Public Health England Variant of Concern) and B.1.351/501Y.V2 (South African Variant of Concern) include multiple mutations in the Spike protein that appear to improve viral infectivity^24-26^ or escape neutralizing antibodies^27,28^. While early data suggested limited impact of the 501Y mutation on neutralizing antibody responses ^29-31^, recent data suggest that this mutation may reduce mRNA vaccine induced neutralization^27^, matching recent Novavax vaccine efficacy results in the UK^32,33^. Conversely, consistently reduced neutralization has been noted against the emerging viral variant from South Africa, in parallel to reduced vaccine efficacy in RSA^27,32^. Yet, how these mutations affect overall vaccine-induced immunity remains unclear. Thus, we profiled the functional antibody response to the UK (B.1.1.7 RBD and N501Y∆69-70 Spike) and RSA (B.1.351 RBD, E484K RBD and E484K Spike) variants compared to the D614G viral variant for spike and WT variant for RBD. Strong correlations were observed between the dominant D614G variant and the two emerging variant viruses, across all antibody metrics (**Fig 6A**). However, some diminution of antibody binding (IgG1/IgG3) and Fc-receptor binding was noted across both strains, albeit the effect was most pronounced for the RSA strains in both the macaques (**Fig 6A**) and humans (**Fig 6B**). Interestingly, macaques (**Fig 6A**) displayed a more pronounced loss of humoral reactivity to the B.1.1.7 and B.1.351 variants compared to humans (**Fig 6B**), where humans with high antibody titers largely retained FcR-binding to the mutants. Conversely, human vaccinees with intermediate antibody titers exhibited a more profound loss of S-specific FcR binding, potentially contributing to the observed vulnerability in a fraction of vaccinees to infection/disease. Multivariate analyses by PCA, where variation in antibody profiles across SARS-CoV-2 variants were captured (**Fig 6C**), point to altered binding, specifically in the macaque humoral immune response across the E484K and B.1.351 RBD variants compared to the UK variant. These data highligh the dominant effect of E484K in knocking out Fc-effector function against the RBD, in addition to a loss of neutralization^27-29,34^ (**Supplementary Fig 2A-C**).

**Figure 6.**
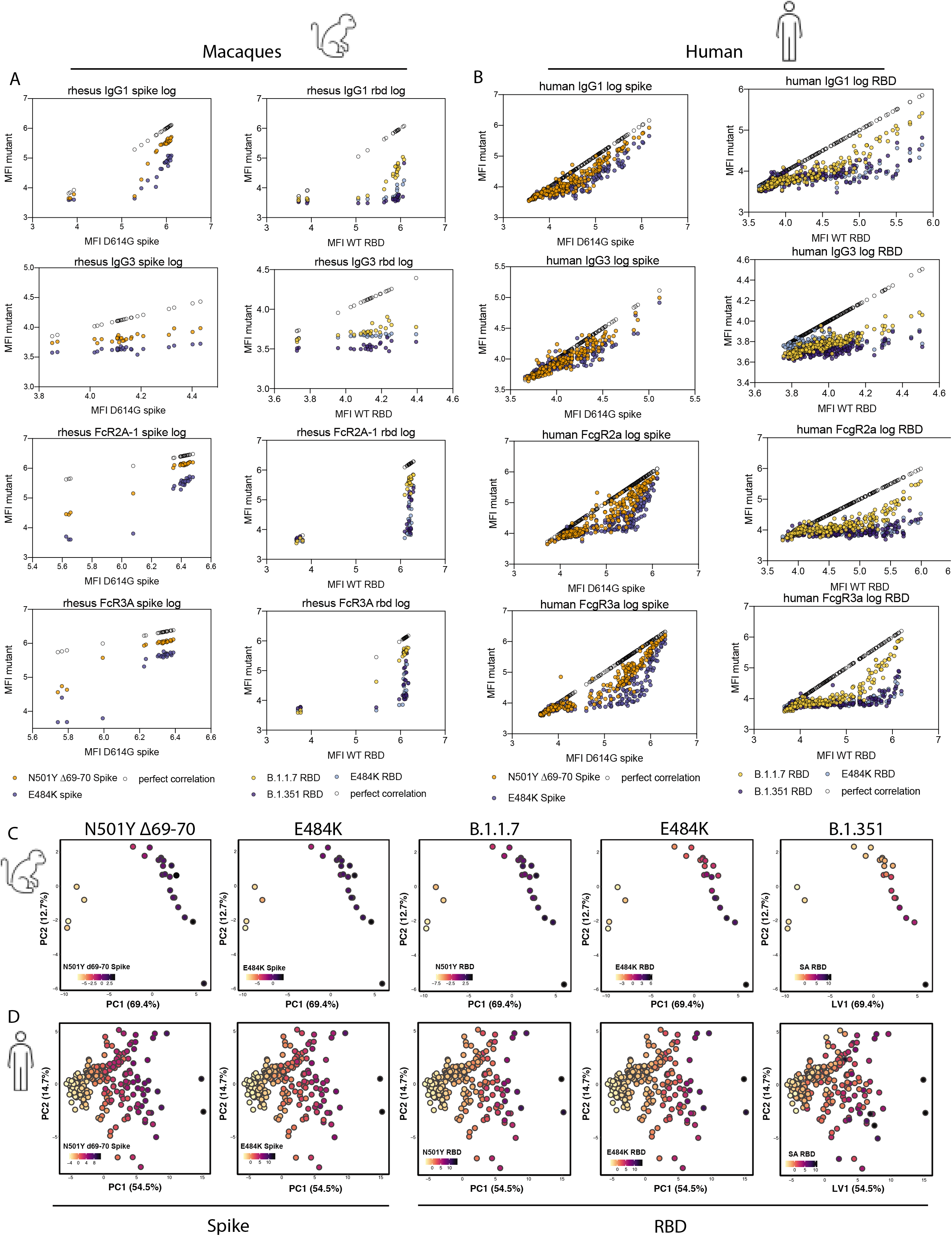
Antibody binding and functionality against emerging SARS-CoV-2 variant spike proteins. (A) NHP Serum (n=24) was collected day 31/32 after the first dose of NVX-CoV2373, and profiled for the antibody response to the emerging SARS-CoV-2 variants, N501Y∆69-70 Spike, E484 Spike, B.1.1.7 RBD, B.1.351 RBD, and E484K RBD. Luminex was used to quantify and correlate the antibody isotypes (IgG1 and IgG3) and FcR binding (FcγRIIA-1 and FcγRIIIA) response between WT Spike protein and SARS-CoV-2 variants Spike or RBD proteins. (B) Serum was collected after vaccinations from individuals (n=237) vaccinated with NVX-CoV2373, and the antibody response was profiled. Luminex was used to quantify and correlated antibody isotypes (IgG1) and FcR binding (FcγRIIA and FcγRIIIA). PCA plots show the multivariate distribution of the antibody profiles across all macaques (C) and humans (D) for Spike (left) or RBD (right) humoral immune responses. The color of the dots represents the sum of the scaled magnitude of the labeled Spike (left) or RBD (right) variant humoral immune response.

These data point to a slight reduction in antibody effector function against the B.1.1.7 variants, but a diminished overall response to the E484K and B.1.351 variant that may explain differences in the level of efficacy observed in the recent clinical trials^32^. The presence of more robust recognition of the full Spike variants among individuals with robust humoral immune responses, representing approximately half of the vaccinees, suggests that at high antibody titers, vaccine induced immunity may contribute to protection against variants, via non-RBD specific responses, the latter that are lost with the E484K mutation. Thus, these data suggest that NVX-CoV2373 stimulates a robust humoral immune response that is fully functionally matured with boosting. While antigen dose has a more limited influence on shaping the functional protective profile of the humoral immune response, the induction of both neutralization and Fc-receptor mediated activity represents key correlates of immunity against upper and lower respiratory tract protection against SARS-CoV-2 and its variants, that may be key to both protection from disease and transmission.

## Discussion

Vaccine shortages, the need for rapid global deployment, increasing reinfection cases, and the emergence of viral variants have collectively pointed to the urgent need to define correlates of immunity against SARS-CoV-2 and its variants. Using a unique vaccine study, poised to profile both the importance of antigen-dose and boosting, here we deeply and comprehensively dissected the key correlates of immunity against upper and lower respiratory tract infection. Despite the induction of robust vaccine-specific antibody titers and neutralization with a single dose or two doses of 5μg or 25μg NVX-CoV2373, differential levels of viral restriction were observed across animals in the upper and lower respiratory tracts. Specifically, animals receiving a single dose vaccine were only partially protected against replicating virus in the upper respiratory tract, whereas animals receiving 2 doses exhibited near complete protection. These data suggest that a single dose may prevent disease, but that two doses may be essential to block further transmission.

The improved protection of the two-dose vaccine was linked to a dramatic maturation of the Fc-effector profiles of vaccine induced antibodies, that collaborated with neutralization as key correlates of immunity against viral replication, with highly functional and neutralizing antibody responses conferring the most robust restriction across the upper and lower respiratory tract. Thus overall, these data demonstrate the critical importance of a coordinated Fab- and Fc-mediated antibody response for full protection against SARS-CoV-2 infection, that may also function against emerging variants.

Both human vaccines, mRNA-1273 and BNT162b2, require a prime and boost to achieve optimal protection. However, as the logistical challenges become apparent in distributing a vaccine globally, interest in increasing the available vaccine by reducing the amount of vaccine or doses given per individual has increased. Preliminary retrospective analysis of the first dose of the Pfizer/BNT162b2 before boosting suggested approximately a 52% protection from severe infection ^5,35^. However, whether a single dose can provide long-term protection remains unclear. While immunogenicity and durability vary significantly across vaccine platforms^5-7,10,11,19^, our data demonstrate some level of protection against lower-respiratory infection after a single vaccine. Yet single dose vaccine-maintained IgM, exhibited incomplete class switching, poor mucosal antibody levels, and demonstrated incomplete functional effector and neutralizing responses, albeit a more balanced response was noted at the higher (25μg) antigen dose. However, after two doses, the explosion of antibody effector function and neutralization likely resulted in a significant increase in protection against both upper and lower respiratory viral replication, linked to the combined presence of potent neutralizing and Fc-effector inducing antibodies and continue to point to the value of the booster immunization.

Neutralizing antibodies represent a critical obstacle to viral infection at the time of infection. However, the density of antibody-producing cells likely varies along the respiratory tract, with a higher density of immune cells found in the lower respiratory tract compared to the more immune barren upper respiratory tract^36,37^. Thus, to achieve complete sterilizing protection from infection in the upper respiratory tract, it is plausible that additional immune mechanisms may be required in the upper respiratory tract to compensate for potentially lower antibody levels. Here we observed the key role of neutralizing antibodies deep within the lungs, but the critical importance of SARS-CoV-2 antibodies of multiple subclasses, binding to multiple Fc-receptors, and complement activation as key additional functional mechanisms that may contribute to upper respiratory protection. Given that the NVX-CoV2373 vaccine induced potent neutralizing antibodies across doses and regimens, we were unable to divorce the influence of neutralization and Fc-effector function. Similar profiles have been noted following reinfection, DNA and Ad26-vaccine studies, marking the co-evolution of the Fab and Fc, and the importance of both ends of the molecule in protective immunity^10,11^. However, whether neutralization and/or Fc-effector function persist differentially over time following vaccination, conferring different levels of protection may provide key insights on precise durable correlates of immunity.

As the virus has begun to adapt to populations across the globe, a number of SARS-CoV-2 variants have begun to emerge. The D641G mutation spread rapidly from Europe to other continents, resulting in a conformational change in the rigidity of the RBD, resulting in enhanced infectivity in vitro, but resulting in no escape from neutralizing antibodies ^38-43^. Similarly, more recently the B.1.1.7 mutation has spread across and out of the UK since September 2020^21^, representing 3 key mutations N501Y, P681H, and ∆69-70, that have been linked to enhanced ACE2-binding, but limited impact on neutralizing antibody activity by monoclonal or the Pfizer/BNT162b2^20,26,28,30,31,44^. Additional variants have begun to emerge in South Africa (B.1.351/501Y.V2) and Brazil (P.1), including mutations both in the RBD and the N-terminal domain of the S-protein, demonstrating significant evasion of antibody-mediated neutralization^28,31,45-49^. Here we noted a loss of both binding and FcR binding activity across the variants in both macaques and humans, with a more profound loss of binding to RSA variant mutations, particularly related to a nearly complete loss of RSA RBD-specific humoral immunity. However, full Spike-specific antibody binding persisted in approximately half of the human vaccinees, particularly in those with robust antibody titers, pointing to the potential for persisting Fc-effector functions as a key compensatory correlate of immunity in the face of evolving mutants that knock out RBD-binding and neutralization. These data mirror the observed rates of protection observed in the recent Phase2 vaccines studies^32^, further substantiating the potential critical importance of both Fab and Fc-functionality in overall population level vaccine efficacy. Yet, further research is needed promptly to identify the impact of emerging mutations on both neutralization and other antibody effector functions that may contribute to antiviral control and protection.

After just 4 months, the WHO declared that the SARS-CoV-2 virus had caused a worldwide pandemic. In response, several vaccines have progressed through late stages of clinical evaluation. To date, messenger RNA (mRNA) vaccines, BNT162b2 and mRNA-1273, recently received Emergency Use Authorization (EUA). Although these vaccines have an acceptable safety profile and effectively protect against more severe disease, they require freezing, have limited data on long-term durability, and have not been shown to protect against infection or transmission. Moreover, given the limited number of vaccine doses available, more vaccine candidates are urgently needed that are able to counteract both wildtype and emerging variant strains. Thus, the need to understand correlates of immunity has never been more urgent, to support the selection and design of additional vaccines able to confer global protective immunity. Here, we describe the identification of correlates of immunity using a subunit vaccine that is stable at refrigerated temperatures, and is immunogenic and well tolerated in human studies^19^. In this study, we demonstrate the presence of binding and neutralizing antibody titers after a single immunization, using either 5μg or 25μg of vaccine, but a remarkable maturation of the Fc-effector profile after a second immunization. Moreover, while partial protection was observed with neutralizing antibodies alone after a single round of immunization, complete protection in the upper and lower respiratory tract was observed with a second round of immunization, marking critical Fab and Fc-mediated correlates of immunity that may be key to both protection against disease and transmission of SARS-CoV-2 and emerging variants. Thus this work bolsters the value of boosting, which will undoubtedly be critical not only to achieve complete protection against infection and transmission, but also to drive durability. Collectively, these data provide key insights into compartment specific immune correlates that may be critical for protection against virus shedding that could help meet an urgent public health need and accelerate the establishment of herd immunity^50,51^.

## Materials and methods

### Cell line, viruses, and receptor

Vero E6 cells were obtained from ATCC, CRL-1586 and maintained in Minimal Eagles Medium (MEM) supplemented with 10% fetal bovine serum (FBS), 1% glutamine, and 1% penicillin and streptomycin (P/S). THP-1 cells (ATCC TIB-202) maintained in Roswell Park Memorial Institute (RPMI) medium, supplemented with 10% FBS, 1% glutamine, 1% P/S, 1% HEPES, and 50µM **β**-ME. HEK 293T/ACE2 cells were obtained from Drs. Michael Farzan and Huihui Mu at the Scripps Research Institute (Jupiter, FL, USA). For the challenge phase of the study, the SARS-CoV-2 (USA-WA-1/2020) passage 4 (P4) isolate was obtained from Biodefense and Emerging Infections Research Resource Repository (catalog number NR-52281, BEI Resources, GenBank accession number MN985325.1). For the in vitro neutralization assay, the SARS-CoV-2 (USA-WA-1/2020) isolate was obtained from the Center for Disease Control and provided by Dr. Matthew Frieman, University of Maryland. Histidine-tagged human ACE2 receptor was purchased from Sino Biologics (Beijing, CHN). Matrix-M™ adjuvant was provided by Novavax, AB (lot number M1-111, Uppsala, SWE)^52^.

### NVX-CoV2373 spike glycoprotein

The SARS-CoV-2 S vaccine was constructed from the full-length, wild-type SARS-CoV-2 S glycoprotein based on the GenBank gene sequence MN908947 nucleotides 21563-25384. The native S protein was modified by mutating the putative furin cleavage site (682-RRAR-685 to 682-QQAQ-685) in the S1/S2 cleavage domain to confer protease resistance. Two additional proline amino acid substitutions were inserted at positions K986P and V987P (2P) within the heptad repeat 1 (HR1) domain to stabilize SARS-CoV-2 S in a prefusion conformation^53^. The synthetic transgene was codon optimized and engineered into the baculovirus vector (BV2373) for expression in *Spodoptera frugiperda* (Sf9) insect cells (GenScript, Piscataway, NJ, USA). NVX-CoV2373 spike trimers were detergent extracted from the plasma membrane with phosphate buffer containing TERGITOL NP-9, clarified by centrifugation, and purified by TMAE anion exchange and lentil lectin affinity chromatography. Purified NVX-CoV2373 (547 μg mL^-1^, lot number BV2373-16APR20) was formulated in 25 mM sodium phosphate (pH 7.2), 300 mM NaCl, and 0.02% (v/v) polysorbate and supplied frozen at −80°C ± 10°C^54^.

### Animal ethics statement

The immunization and challenge phases of the study complied with all applicable sections of the Final Rules of the Animal Welfare Act regulations (9 CFR Parts 1, 2, and 3) and *Guide for the Care and Use of Laboratory Animals - National Academy Press, Washington D. C. 8th Edition, 2011 (The Guide*). The study was conducted at the Texas Biomedical Research Institute (Texas Biomed, San Antonio, TX, USA), an AAALAC (Association for the Assessment and Accreditation of Laboratory Animal Care) accredited facility. The work was conducted in accordance with a protocol approved by Texas Biomed’s Institutional Animal Care and Use Committee.

### Human ethics statement

The Phase 1 vaccine study was previously described^19^. Healthy 18-59-year-old men and non-pregnant women were included in the study. Previously infected individuals were excluded. With the exception of 6 sentinel participants vaccinated in an open-label manner, the remaining 125 participants were randomly assigned to vaccine and placebo groups in a blinded fashion. All subjects signed informed consent and safety oversight was monitored by a data monitoring board.

### Animal husbandry

Animals were housed individually in stainless steel cages with wire mesh bottoms. Animals were fed commercially available certified primate diet from Purina Mills 5048 (LabDiet) and provided water *ad libitum* from an institutional watering system that was analyzed monthly for impurities. Environmental conditions included 12 hour light and 12 hour dark cycle with controlled temperature (74°F ± 10°F) and humidity (30% to 70% RH). Cages were cleaned daily.

Twenty-four experimentally naïve rhesus macaques (*Macaca mulatta*) of Chinese origin were sourced from Envigo (Alice, TX, USA). Animals were screened and determined to be negative for Simian Immunodeficiency Virus (SIV), Simian T-Lymphotropic Virus-1 (STLV-1), Simian Varicella Virus (SVV) and *Macacine herpesvirus* 1 (Herpes B virus), and Simian Retrovirus (SRV1 and SRV2) by polymerase chain reaction (PCR), and negative for *Trypanosoma cruzi*. Rectal swabs were collected and tested for Shigella, Campylobacter, Salmonella, and Yersinia. Pharyngeal swabs were used to test for *Bordetella bronchiseptica*. All animals were tested and verified to be negative for tuberculosis.

The vaccination phase of the study was performed in the Texas Biomed Animal Biosafety Level 2 (ABSL-2) facility. Following the immunization phase of the study, animals were transferred and acclimated for 7 days in the Texas Biomed ABSL-3 facility prior to challenge. Animals were monitored a minimum of twice daily for the duration of the study.

### Study blinding

This study was blinded (assignment to vaccinated/immunized versus placebo group) to avoid bias in evaluation, euthanasia, gross pathology assessment, and qRT-PCR assay outcome. All staff performing in vitro assays were blinded to the animal vaccine dosage and to whether the animal received vaccine or placebo while performing assays and analysis.

### Study design

Animals were randomly assigned to groups, with stratification across age and gender, using a computerized randomization procedure. Twenty-four (12 male and 12 female) rhesus macaques, within the age range of >3 to <8-year-olds and weight range ≥3.67 kg to ≤10 kg, were randomized into four immunization groups and two placebo groups. NVX-CoV2373 was formulated with 50μg Matrix-M on the day of immunization. The placebo groups received formulation buffer. Groups 1 (1 male and 1 female) received placebo in two doses spaced 21 days apart (study day 0 and 21) and group 4 (1 male and 1 female) received placebo in one dose (study day 0). Group 2 (2 females and 3 males) received 5μg NVX-CoV2373 + 50μg Matrix-M and group 3 (2 females and 3 males) received 25μg NVX-CoV2373 + 50μg Matrix-M in two doses spaced 21 days apart (study day 0 and 21). Group 5 (3 females and 2 males) received 5μg NVX-CoV2373 + 50μg Matrix-M and group 6 (3 females and 2 males) received 25μg NVX-CoV2373 + 50μg Matrix-M in one dose (study day 0). Injections (0.5 mL) were administered in the thigh muscle.

Animals were sedated by intramuscular (IM) administration of Telazol (2-8 mg kg^-1^, IM) prior to vaccination, collection of blood samples, virus challenge, collection of nasal swabs, nasal washes, and bronchoalveolar lavage (BAL). For serologic assessments, serum was collected on study day 0 prior to immunization and day 21, and day 31 or 32 after the first immunization and stored at −80°C until assayed. Nasal washes, nasal pharyngeal swabs, and BAL were collected on study day 31/32, prior to challenge.

### Anti-spike IgG and IgA ELISA

Serum, nasal wash, and BAL anti-SARS-CoV-2 spike (S) protein IgG titers were determined by ELISA. Briefly, 96-well plates (Thermo Fisher Scientific, Rochester, NY, USA) were coated with 1.0 µg mL^-1^ of SARS-CoV-2 S protein (BV2373, Lot# 16Apr20, Novavax, Inc. Gaithersburg, MD, USA). Plates were washed with phosphate buffered Tween (PBS-T) and non-specific binding was blocked with TBS Startblock blocking buffer (Thermo Fisher Scientific, Rochester, NY, USA). Serum samples were serially diluted 3-fold starting with a 1:100 dilution and BAL and nasal wash samples were serially diluted 2-fold starting with a 1:2 dilution, then added to the coated plates and incubated at room temperature for 2 hours. For IgG ELISA, plates were washed with PBS-T, then incubated with horseradish peroxidase (HRP)-conjugated mouse anti-monkey IgG (catalog number 4700-05, Southern Biotech, Birmingham, AL, USA) for 1 hour. For IgA ELISA, plates were washed with PBS-T and mouse anti-monkey IgA (catalog number MCA2553, Bio-Rad, Hercules, CA, USA) was added for 1 hour followed by washing with PBS-T, then incubation with HRP-conjugated goat anti-mouse IgG (catalog number 1030-05, Southern Biotech). Plates were then developed with 3,3’,5,5’-tetramethylbenzidine (TMB) peroxidase substrate (Sigma, St. Louis, MO, USA). Reactions were stopped with TMB stop solution (ScyTek Laboratories, Inc. Logan, UT, USA). Plates were read at OD 450 nm with a SpectraMax Plus plate reader (Molecular Devices, Sunnyvale, CA, USA). EC_50_ values and endpoint titer values were calculated by 4-parameter fitting using SoftMax Pro 6.5.1 GxP software. Individual animal anti-S IgG or IgA titers, and group geometric mean titer (GMT) and 95% confidence interval (95% CI) were plotted using GraphPad Prism 9.0 software. For serum titers below the assay lower limit of detection (LOD), a titer of < 1000 (starting dilution) was reported and a value of “50” assigned to the sample to calculate the group mean titer. For BAL and nasal wash titers below the assay LOD, a titer of <2 (starting dilution) was reported and a value of “1” assigned to the sample to calculate the group mean titer.

### Human angiotensin converting enzyme 2 (hACE2) receptor blocking antibody

Human ACE2 receptor blocking antibody titer was determined by ELISA. Ninety-six well plates were coated with 1.0 μg mL^-1^ SARS-CoV-2 rS protein (BV2373, lot no. 16Apr20, Novavax, Inc., Gaithersburg, MD, USA) overnight at 4°C. Sera were serially diluted 2-fold starting with a 1:20 dilution and were added to coated wells for 1 hour at room temperature. After washing, 30 ng mL^-1^ histidine-tagged hACE2 (Sino Biologics, Beijing, CHN) was added to wells for 1 hour at room temperature. HRP-conjugated mouse anti-histidine-tag IgG (1:4000) (catalog number 4603-05, Southern Biotech, Birmingham, AL, USA) was added for 1 hour followed by addition of TMB substrate. Plates were read at OD 450 nm with a SpectraMax Plus plate reader (Molecular Devices, Sunnyvale, CA, USA) and data analyzed with SoftMax Pro 6.5.1 GxP software. The % Inhibition for each dilution for each sample was calculated using the following equation in the SoftMax Pro program: 100-[(MeanResults/ControlValue@PositiveControl)∗100].

Serum dilution versus % Inhibition plot was generated, and curve fitting was performed by 4-parameter logistic (4PL) curve fitting to data. Serum antibody titer at 50% inhibition (IC_50_) of hACE2 to SARS-CoV-2 S protein was determined in the SoftMax Pro program. The group GMT and 95% CI and individual animal titers were plotted using GraphPad Prism 9.0 software. For a titer below the assay lower limit of detection (LOD), a titer of < 20 (starting dilution) was reported and a value of “10” assigned to the sample to calculate the group mean titer.

### SARS-CoV-2 neutralizing antibody assay

The SARS-CoV-2 neutralizing antibody assay was conducted in a select agent ABSL-3 containment facility at the University of Maryland, School of Medicine. Sera were diluted 1:20 in Vero E6 cell growth media and further serially diluted 1:2 to 1:40,960. SARS-CoV-2 (multiplicity of infection (MOI) of 0.01 pfu per cell) was added to the wells for 60 min at 37°C. Vero E6 media was used as a negative control. Each serum dilution was assessed microscopically for inhibition of virus cytopathic effect (CPE) on Vero E6 cells. The endpoint titer was reported as the reciprocal of the dilution at which 99% CPE was observed at 3 days post infection^19,54^.

### Preparation of the SARS-CoV-2 challenge stock

A fourth cell-culture passage (P4) of SARS-CoV-2 isolate USA-WA1/2020 was obtained from Biodefense and Emerging Infections Research Resources Repository (catalog number NR-52281, BEI Resources, GenBank accession number MN985325.1). Live virus stock was prepared in the Texas Biomed ABSL-3 containment facility. The stock virus was passaged for a fifth time (P5) in Vero E6 cells at a MOI of 0.001 to produce the master virus stock. The master stock was again passaged in Vero E6 cells at a MOI of 0.02 (P6) to produce the challenge stock. The P6 challenge stock had a titer of 2.10 × 10^6^ pfu mL^-1^ (Lot No. 20200320) and was stored 500 μL aliquots at −65°C in Dulbecco’s modified essential media (DMEM) and 10% fetal bovine serum. The identity of the challenge stock was confirmed to be SARS-CoV-2 by deep sequencing and was confirmed to be identical to the published sequence (GenBank: MN985325).

### SARS-CoV-2 challenge

Vaccinated and placebo animals were transferred from the ABSL-2 facility on study day 31/32 to the ABSL-3 facility and acclimated for 7 days. On the day of challenge (study day 38), animals were sedated and challenged with a total target dose of 1.05 × 10^6^ pfu in 500 μL. The challenge dose was equally administered by the intranasal (IN) route 5.25 × 10^5^ pfu in 250 μL and intra-tracheal (IT) route 5.25 × 10^5^ pfu in 250 μL. IN administration was performed with an atomization device (Teleflex Intranasal Mucosal Atomization Device LMA MAD Nasal Device, Morrisville, NC, USA) and IT delivery was performed with Tracheal Mucosal Atomization Device (Teleflex Laryngo-Tracheal Mucosal Atomization Device LMA MADGIC, Morrisville, NC, USA). To confirm the challenge dose, aliquots of the challenge samples were collected prior to challenging the first animal and last animal and stored at ≤ −65°C. A neutral red agarose overlay and conventional plaque assay were used to confirm the titer of the challenge dose. The actual pre- and post-challenge titers were 1.80 × 10^6^ pfu mL^-1^ and 7.83 × 10^5^ pfu, respectively.

### Sample collection for SARS-CoV-2 RNA quantification

#### Nasal pharyngeal swab collection

Animals were sedated and nasal pharyngeal swabs were collected prior to challenge (study day 31/32) and on 2, 3, 4, 6, and 7-8 days post infection (dpi). After collection, swabs were placed in a tube containing viral transport medium (VTM), then stored at ≤-60°C until processing.

#### Bronchoalveolar lavage (BAL) collection

BAL aspirates were collected prior to challenge (study day 31/32) and on 2, 4 and 7-8 dpi. Animals were sedated and the trachea visualized with a laryngoscope. A sterile rubber feeding tube with stylet was inserted into the trachea and into the airway until it met slight resistance. Up to 80 mL of warm (<40°C) sterile saline, divided into multiple aliquots, was instilled through the tube. Aspirated fluid was dispensed into sterile vials with VTM and stored at ≤-60°C until batch processed.

#### Nasal wash collection

Nasal washes were collected prior to infection (study day 31/32) and 2, 4, and 7-8 dpi. Animals were sedated and a syringe with a flexible tipped 20-22-gauge intravenous (IV) catheter was inserted into the nostril passage and a volume of 2.5-5mL of sterile saline instilled. Samples were collected in sterile conical tubes containing VTM and stored at ≤-60°C until batch processed.

#### Tissue collection

Tissues were collected 7-8 dpi (study days 45-46) at the scheduled necropsy from the upper, middle and lower lobes of the lung; nasal cavity; and trachea. Tissues were weighed and stored at 80°C ± 10°C until batch processed. RNA was extracted analyzed for the presence of SARS-CoV-2 RNA via qRT-PCR targeting the N1 gene.

### Quantification of virus load in nasal swabs/washes, BAL, and tissues

#### Genomic (g)RNA virus

Samples were assessed for viral load by qRT-PCR. A 250 µL aliquot of VTM inactivated in TRIzol LS reagent (catalog number 10296010, ThermoFisher Scientific) was used for isolation of total RNA. For total viral RNA, qRT-PCR targeting the nucleocapsid gene (N1) was run on duplicate samples and results reported as genome equivalents (GE) mL^-1^ for nasal washes/swabs and BAL. For tissue samples, results are reported as GE μg^-1^ for tissue homogenates.

#### Subgenomic (sg)RNA virus

Replicating virus load by qRT-PCR targeting the subgenomic envelope (E) gene RNA in 250 µL aliquot of nasal swabs, nasal washes, and BAL aspirates. The forward and reverse primers, probe, cycling conditions, and Master Mix included:

SUBGEN-FORWARD: CGATCTCTTGTAGATCTGTTCTC

E_Sarbeco_R2 Reverse Primer: ATATTGCAGCAGTACGCACACA

Probe (Thermo): FAM-MGB: ACACTAGCCATCCTTACTGCGCTTCG

TaqPath™ 1-Step RT-qPCR Master Mix, CG (catalog number A15299, ThermoFisher Scientific). Cycling parameters were 25°C 2 minutes, 50°C 15 minutes, 95°C 2 minutes; Amplification 40 × 95°C 3 seconds, 60°C 30 seconds.

### Pseudovirus neutralizing antibody assay

SARS-CoV-2 neutralization was assessed with spike-pseudotyped virus infection of HEK 293T/ACE2 cells as a function of reduction in luciferase (Luc) reporter activity. HEK 293T/ACE2 cells were maintained in DMEM containing 10% fetal bovine serum, 25 mM HEPES, 50 µg mL^-1^ gentamycin and 3 µg mL^-1^ puromycin. An expression plasmid encoding codon-optimized full-length spike of the Wuhan-1 strain (VRC7480), was provided by Drs. Barney Graham and Kizzmekia Corbett at the Vaccine Research Center, National Institutes of Health (USA). The D614G amino acid change was introduced into VRC7480 by site-directed mutagenesis using the QuikChange Lightning Site-Directed Mutagenesis Kit (catalog number 210518, Agilent Technologies). The mutation was confirmed by full-length spike gene sequencing. Pseudovirions were produced in HEK 293T/17 cells (ATCC cat. no. CRL-11268, Manassas, VA, USA) by transfection using Fugene 6 (catalog number E2692, Promega, Madison, WI, USA) and a combination of spike plasmid, lentiviral backbone plasmid (pCMV ΔR8.2) and firefly Luc reporter gene plasmid (pHR’ CMV Luc) in a 1:17:17 ratio^55^. Transfections were allowed to proceed for 16-20 hours at 37°C. Medium was removed, monolayers rinsed with growth medium, and 15 mL of fresh growth medium added. Pseudovirus-containing culture medium was collected after an additional 2 days of incubation and was clarified of cells by low-speed centrifugation and 0.45µm micron filtration and stored in aliquots at −80°C. TCID_50_ assays were performed on thawed aliquots to determine the infectious dose for neutralization assays (RLU 500-1000x background, background 50-100 RLU).

For neutralization, a pre-titrated dose of virus was incubated with 8 serial 5-fold dilutions of serum samples in duplicate in a total volume of 150 µL for 1 h at 37°C in 96-well flat-bottom poly-L-lysine-coated Biocoat plates (catalog number 354461, Corning, NY, USA). Cells were suspended using TrypLE Select Enzyme solution (Thermo Fisher Scientific) and immediately added to all wells (10,000 cells in 100 µL of growth medium per well). One set of 8 control wells received cells + virus (virus control) and another set of 8 wells received cells only (background control). After 66-72 h of incubation, medium was removed by gentle aspiration and 30 µL of Promega 1X lysis buffer was added to all wells. After a 10 minute incubation at room temperature, 100 µL of Bright-Glo luciferase reagent was added to all wells. After 1-2 minutes, 110 µL of the cell lysate was transferred to a black/white plate (Perkin-Elmer). Luminescence was measured using a PerkinElmer Life Sciences, Model Victor2 luminometer. Neutralization titers are the serum dilution at which relative luminescence units (RLU) were reduced by either 50% (ID_50_) compared to virus control wells after subtraction of background RLUs. Serum samples were heat-inactivated for 30 min at 56°C prior to assay.

### Antibody-dependent cellular phagocytosis and neutrophil phagocytosis

ADCP and ADNP were conducted as previously described^56,57^. Briefly, NVX-CoV2373 Spike protein was biotinylated using EDC (Thermo Fisher) and Sulfo-NHS (Thermo Fisher), and then coupled to yellow/green Neutravidin-conjugated beads (Thermo Fisher). Immune complexes were formed by incubating the bead+protein conjugates with diluted serum for 2 hours at 37°C, and then washed to remove unbound antibody. The immune complexes were then incubated overnight with THP-1 cells (ADCP), or for 1 hour with RBC-lyzed whole blood (ADNP). THP-1 cells were then washed and fixed in 4% PFA, while the RBC-lyzed whole blood was washed, stained for CD66b+ (Biolegend) to identify neutrophils, and then fixed in 4% PFA. Flow cytometry was performed to identify the percentage of quantity of beads phagocytosed by THP-1 cells or neutrophils, and a phagocytosis score was calculated (% cells positive × Median Fluorescent Intensity of positive cells). Flow cytometry was performed with an IQue (Intellicyt) or LSRII(BD) and analysis was performed using IntelliCyt ForeCyt (v8.1) or FlowJo V10.7.1.

### Antibody-dependent complement deposition

ADCD was conducted as previously described^58^. Briefly, NVX-CoV2373 Spike protein was biotinylated using EDC (Thermo Fisher) and Sulfo-NHS (Thermo Fisher), and then coupled to red Neutravidin-conjugated microspheres (Thermo Fisher) or directly coupled to Carboxylate-Modified microspheres (Thermo Fisher). Immune complexes were formed by incubating the bead+protein conjugates with diluted serum for 2 hours at 37°C, and then washed to remove unbound antibody. The immune complexes were then incubated with lyophilized guinea pig complement (Cedarlane) and diluted in gelatin veronal buffer with calcium and magnesium (Boston Bioproducts) for 30 minutes. C3 bound to immune complexes was detected by fluorescein-conjugated goat IgG fraction to guinea pig Complement Ce (MP Biomedicals). Flow cytometry was performed to identify the percentage of beads with bound C3. Flow cytometry was performed with an IQue (Intellicyt) and analysis was performed on IntelliCyt ForeCyt (v8.1).

### Antibody-dependent NK cell degranulation

Antibody-dependent NK cell degranulation was conducted as previously described^59^. NVX-CoV2373 Spike protein was coated on Maxisorp ELISA plate (Thermo Fisher), and then blocked with 5% BSA. Antibodies were then added and incubated for 2 hours at 37°C. Human NK cells were isolated from peripheral blood by negative selection using the RosetteSep Human NK cell enrichment cocktail following the manufacturer’s instructions. Human NK cells were then added to the bound antibody and incubated for 5 hours at 37°C in the presence of RMPI+10%FBS, GolgiStop (BD), Brefeldin A (Sigma), and anti-human CD107a antibody (BD Bioscience). After incubation, cells were washed, stained with CD16, CD56, and CD3 (BD Bioscience), and fixed in 4% PFA for 15 minutes. Intracellular staining was performed using the FIX/PERM Cell fixation and permeabilization kit (Thermo), and cells were stained for interferon-γ and macrophage inflammatory protein-1β (BD bioscience). Flow cytometry was performed with an IQue (Intellicyt) and analysis was performed on IntelliCyt ForeCyt (v8.1).

### Isotype and FcR-binding Luminex profiling

Isotyping and FcR profiling was conducted as previously described^60,61^. Briefly, antigens (NVX-CoV2373 Spike, SARS-CoV-2 Spike, S1, RBD, S2, HKU-1 RBD, or OC43 RBD) were carboxyl coupled to magnetic Luminex microplex carboxylated beads (Luminex Corporation) using NHS-ester linkages with Sulfo-NHS and EDC (Thermo Fisher), and then incubated with serum for 2 hours at 37°C. Isotyping was performed by incubating the immune complexes with secondary mouse-anti-rhesus antibody detectors for each isotype (IgG1, IgG2, IgG3, IgG4, IgA), then detected with tertiary anti-mouse-IgG antibodies conjugated to PE. FcR binding was quantified by incubating immune complexes with biotinylated FcRs (FcγR2A-1, FcγR2A-2, FcγR3A, courtesy of Duke Protein Production Facility) conjugated to Steptavidin-PE (Prozyme). Flow cytometry was performed with an IQue (Intellicyt) and analysis was performed on IntelliCyt ForeCyt (v8.1).

### Multivariate analysis

A principal component analysis (PCA) was performed based on serological features using the R package ‘ropls’. The systems serology antibody titers, FcR binding and ADCD measurements were log10-transformed, and all measurements were z-scored. The PCA analyses was performed in R version 4.0.2.

### Statistical analysis

Statistical analyses were performed with GraphPad Prism 9.0 software. Serum antibodies were plotted for individual animals and the geometric mean titer (GMT) and 95% confidence intervals plotted. Virus loads were plotted as the median value, interquartile range, and minimum and maximum values. Student’s t-test or two-way ANOVA was used to determine differences between paired groups as indicated in the figure legends. p ≤ 0.05 was considered significant. The AUCs and bootstrap confidence intervals were calculated using the R package ‘pROC’. For the case of AUC = 1 no confidence interval was provided. To visualize fold change of mutant humoral features with respect to the WT, a volcano plot was constructed. To calculate p-values, the R package ‘stats’ was used. The AUC and fold change analyses were performed in R version 4.0.2.

## Supporting information

Raw Data

## Funding statement and Acknowledgements

This work was funded by Operation Warp Speed. We thank Colin Mann and Kathryn Hastie for production of Spike antigens. We thank Nancy Zimmerman, Mark and Lisa Schwartz, an anonymous donor (financial support), Terry and Susan Ragon, and the SAMANA Kay MGH Research Scholars award for their support. We acknowledge support from the Ragon Institute of MGH, MIT and Harvard, the Massachusetts Consortium on Pathogen Readiness (MassCPR), the NIH (3R37AI080289-11S1, R01AI146785, U19AI42790-01, U19AI135995-02, U19AI42790-01, 1U01CA260476 – 01, CIVIC75N93019C00052), the Gates foundation Global Health Vaccine Accelerator Platform funding (OPP1146996 and INV-001650), and the Musk Foundation.

## Author contributions

NP, MGX, YG, RC, JHT, BZ, MJM, ADP, MJG, CA, AZ, GA, CL, KMP, EOS, DL, FK, and GS contributed to conceptualization of experiments, generation of data and analysis, and interpretation of the results. NP, JHT, BZ, SM, YG, RC, CA, MJG, AZ, DY, KB, FA, SS, SM, MEM, JL, CM, and MBF performed experiments. NP, MGX, YG, RC, GA, and MBF coordinated projects. GS, GG, DL, DM, MGX, AMG, NP, YG, RC, MBF, MJG, CA, GA, CL, KMP, and LE contributed to drafting and making critical revisions with the assistance of others.

## Figure Legends

**Supplementary Figure 1. Gating strategy for flow cytometry**

Example of flow cytometry gating scheme for (A) antibody-dependent cellular phagocytosis (ADCP), (B) antibody-dependent neutrophil phagocytosis (ADNP), (C) antibody-dependent complement deposition (ADCD), and (D) antibody-dependent NK degranulation (measured by CD107%) (NKdegran).

**Supplementary Figure 2. Antibody and FcR binding against emerging SARS-CoV-2 variant spike proteins**

(A) Loadings plot for PC 2 of the NHP PCA model in Figure 6. Features are colored by variant protein. (B) Loadings plot for PC 2 of the Human PCA model in Figure 6. Features are colored by variant protein. (C) Volcano plot of each humoral feature for the immunized NHPs. The fold change in respect to the D614G Spike for spike features or WT RBD for RBD features is plotted in the volcano plot. (B) Volcano plot of each humoral feature for the vaccinated individuals. The fold change in respect to the D614G Spike for spike features or WT RBD for RBD features is plotted in the volcano plot.

