## Supplementary figures and images for "Collaboration between the Fab and Fc contribute to maximal protection against SARS-CoV-2 in nonhuman primates following NVX-CoV2373 subunit vaccine with Matrix-M™ vaccination"

### Supplemental Figure 1

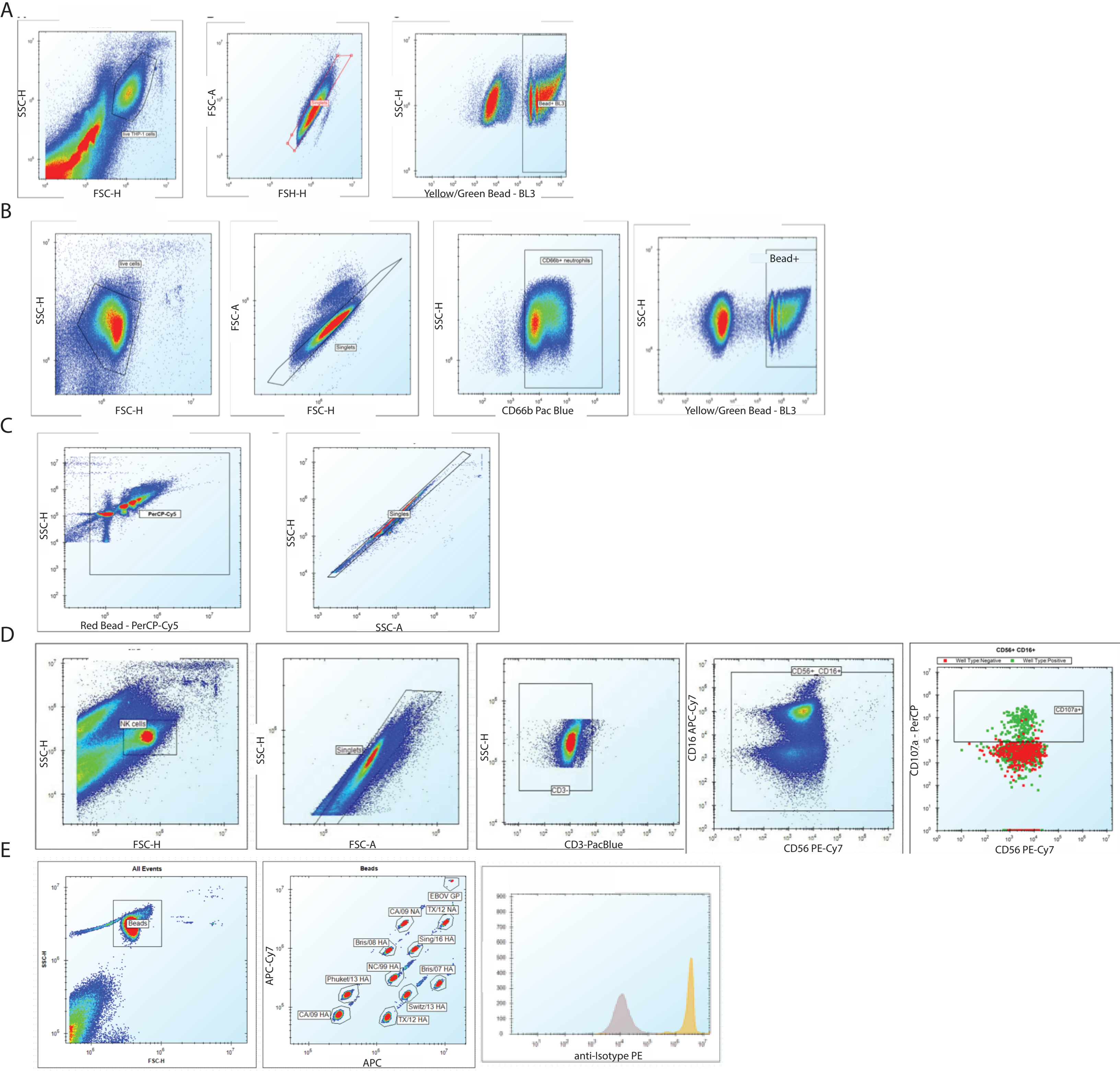

### Supplemental Figure 2

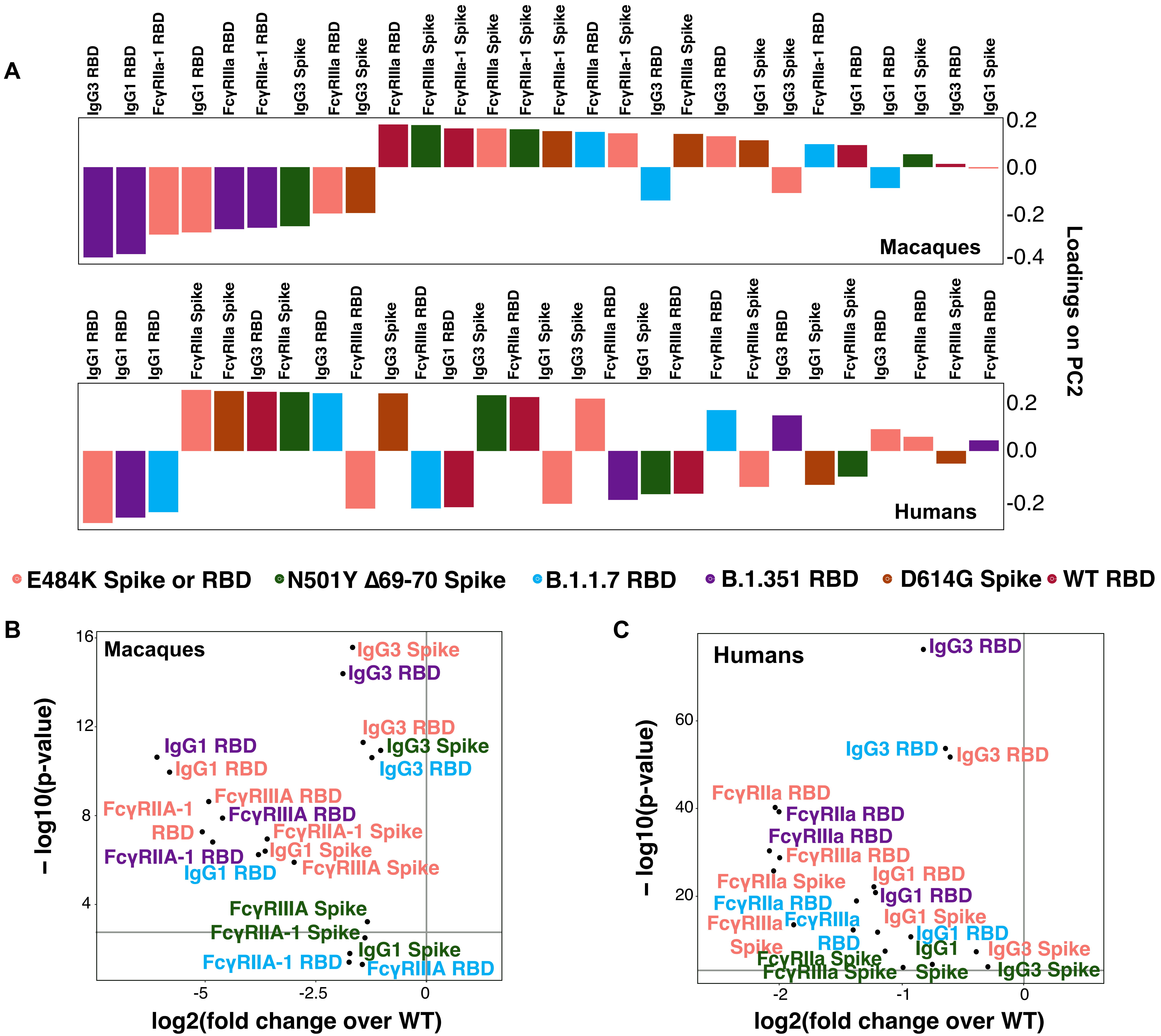
